# Hyperinsulinemia promotes aberrant histone acetylation in triple negative breast cancer

**DOI:** 10.1101/503201

**Authors:** Parijat Senapati, Christine Thai, Angelica Sanchez, Emily J Gallagher, Derek LeRoith, Victoria L. Seewaldt, David K. Ann, Dustin E. Schones

**Author notes:** Corresponding Author: Dustin Schones, Ph.D., Department of Diabetes Complications and Metabolism, City of Hope, Duarte, CA, 91010.

## Abstract

Excess levels of insulin relative to glucose in the blood, or hyperinsulinemia, is considered to be a poor prognostic indicator for patients with triple negative breast cancer (TNBC). While this association has been recognized for some time, the mechanistic role of hyperinsulinemia in promoting TNBC remains unclear. We show that insulin treatment leads to genome-wide increase in histone acetylation, in particular at H3K9, through the PI3K/AKT/mTOR pathway in MDA-MB-231 cells. Genome-wide analysis showed that the increase in histone acetylation occurs primarily at gene promoters. In addition, insulin induces higher levels of reactive oxygen species and DNA damage foci in cells. *In vivo*, hyperinsulinemia also enhances growth of MDA-MB-231 derived tumors through increased histone acetylation. These results demonstrate the impact of hyperinsulinemia on altered gene regulation through chromatin and the importance of targeting hyperinsulinemia-induced processes that lead to chromatin dysfunction in TNBC.

## Introduction

Triple negative breast cancer (TNBC) is a clinically aggressive subtype of breast cancer that does not express estrogen receptor (ER), progesterone receptor (PR) or human epidermal growth factor receptor 2 (HER2) (Carey, Winer, Viale, Cameron, & Gianni, 2010). Recent epidemiological studies suggest that metabolic syndrome and associated disorders are additional risk factors for developing TNBC in premenopausal women (Pierobon & Frankenfeld, 2013). Multiple factors common to metabolic syndrome, including hyperinsulinemia, hyperglycemia, hyperlipidemia, altered adiponectin and leptin levels can contribute to promote tumor growth and progression (Hursting et al., 2012). However, it has been shown that hyperinsulinemia alone even in the absence of obesity and T2D is also associated with increased breast cancer incidence (Del Giudice et al., 1998; Gunter et al., 2009; Lawlor, Smith, & Ebrahim, 2004; Lipscombe, Goodwin, Zinman, McLaughlin, & Hux, 2006) and adverse prognosis (Goodwin et al., 2002; Goodwin et al., 2012).

Hyperinsulinemia can influence tumor growth by multiple mechanisms. Insulin can stimulate tumor cell survival and proliferation by signaling through the insulin receptor (IR) (Belfiore, Frasca, Pandini, Sciacca, & Vigneri, 2009; Huang et al., 2011) as well as by enhancing the available pool of insulin-like growth factor (IGF-I) by decreasing the expression of IGF binding protein (IGFBP1) (Calle & Kaaks, 2004). Increased IR expression and the presence of phosphorylated IR/IGF-IR in breast cancer are associated with poor prognosis and decreased survival (Belfiore et al., 1996; Mathieu, Clark, Allred, Goldfine, & Vigneri, 1997). Interestingly, in a mouse model of hyperinsulinemia without confounding factors such as obesity, hyperglycemia or hyperlipidemia (Novosyadlyy et al., 2010), endogenous hyperinsulinemia increases mammary tumor growth as well as metastases (Ferguson et al., 2012) by signaling primarily through the IR (Gallagher et al., 2013; Novosyadlyy et al., 2010).

Insulin binding to IR leads to downstream activation of PI3K-AKT and MAPK signaling pathways (Saltiel & Kahn, 2001). The PI3K-AKT pathway has oncogenic activity and induces mTOR signaling that promotes cell growth (Engelman, Luo, & Cantley, 2006; Manning & Cantley, 2007). Breast cancers frequently show deregulation of the PI3K-AKT and mTOR pathway (Campbell et al., 2004; Guertin & Sabatini, 2007; Hynes & Boulay, 2006; Vivanco & Sawyers, 2002). Moreover, activation of AKT/mTOR signaling is associated with poor prognosis in TNBCs (Ueng et al., 2012). Since insulin can activate the AKT/mTOR pathway, it is predicted that hyperinsulinemia may drive the aggressive biology of TNBCs in women with insulin resistance and TNBCs. Additionally, mTOR signaling stimulates mitochondrial biogenesis and activity thereby enhancing mitochondrial processes such as TCA cycle, oxidative phosphorylation and ATP production (Morita et al., 2013). Cancer cells exhibit enhanced utilization of glucose which is further converted to citrate by the TCA cycle. In proliferating cancer cells, citrate is converted to acetyl-coenzyme A (acetyl-CoA) that is utilized for production of lipids (Vander Heiden, Cantley, & Thompson, 2009). Acetyl-CoA is also utilized by nuclear histone acetyltransferases as a substrate for histone acetylation, which is an important mechanism for regulating cellular gene expression by aiding in the accessibility of chromatin through several mechanisms (Di Cerbo & Schneider, 2013; Su, Wellen, & Rabinowitz, 2016). Cancer cells often exhibit deregulation of histone acetylation (Di Cerbo & Schneider, 2013). Although several studies indicate the role of insulin signaling in breast cancer, the mechanistic details of these cellular processes in promoting TNBCs is lacking. Most importantly, the effect of hyperinsulinemia on chromatin and gene expression in the nucleus is unclear.

We report here an investigation into the impact of hyperinsulinemia on chromatin acetylation by profiling histone H3 acetylation at lysine 9 (H3K9ac) after insulin treatment of the triple negative breast cancer cell line MDA-MB-231. We show that insulin induces global increases in histone acetylation through the PI3K/AKT/mTOR pathway. We used quantitative ChIP-seq analyses (ChIP-Rx) to show that gene promoters exhibit the greatest increases in histone acetylation. In addition, insulin induces DNA damage in cells through enhanced reactive oxygen species (ROS) production and increased chromatin accessibility. Moreover, increased insulin levels enhance growth of MDA-MB-231 xenograft tumors through enhanced histone acetylation. These findings suggest that hyperinsulinemia leads to altered cell metabolism that influences chromatin acetylation in tumor cells, thereby influencing gene expression across the genome.

## Results

### Insulin induces genome-wide histone acetylation in MDA-MB-231 cells

To investigate the effect of insulin on chromatin, we treated MDA-MB-231 cells with 100nM insulin for different durations and assayed the levels of histone acetylation via western blot. We observed an increase in total histone H3 acetylation levels (acH3) after 3h of insulin treatment (Figure 1A). This increase was more pronounced for specific residues such as H3K9 and H3K14 (Figure 1A). To confirm that the PI3K-AKT pathway was activated by insulin, we further probed the blots with phospho-AKT antibody. Phospho-AKT levels were induced at higher levels after 1h of insulin treatment, after which the signal was attenuated, as expected (Figure 1A). To assess the role of PI3K-AKT/mTOR pathway activation in increasing histone acetylation levels, we used inhibitors targeting mTOR and PI3K kinases. We pre-treated MDA-MB-231 cells with mTOR inhibitor rapamycin and PI3K inhibitor LY294002 followed by insulin treatment. Insulin induced phosphorylation of AKT and p70 S6 kinase (S6K), an mTOR kinase substrate, in rapamycin untreated cells (Figure 1B, lanes 1 and 2). Rapamycin pre-treatment, however, inhibited the phosphorylation of S6K without affecting AKT phosphorylation confirming that rapamycin indeed inhibited mTOR kinase activity (Figure 1B, lanes 3 and 4). Both mTOR inhibition and PI3K inhibition, by rapamycin and LY294002 treatment respectively, inhibited the H3K9ac increase induced by insulin (Figure 1B and 1C). To confirm that the insulin induced histone H3 acetylation is chromatin-bound and not on newly synthesized or free histones, we performed chromatin fractionation after insulin treatment followed by western blot analyses. Results showed that insulin induced H3K9ac was exclusively chromatin bound (Figure S1A-B).

**Figure 1.**
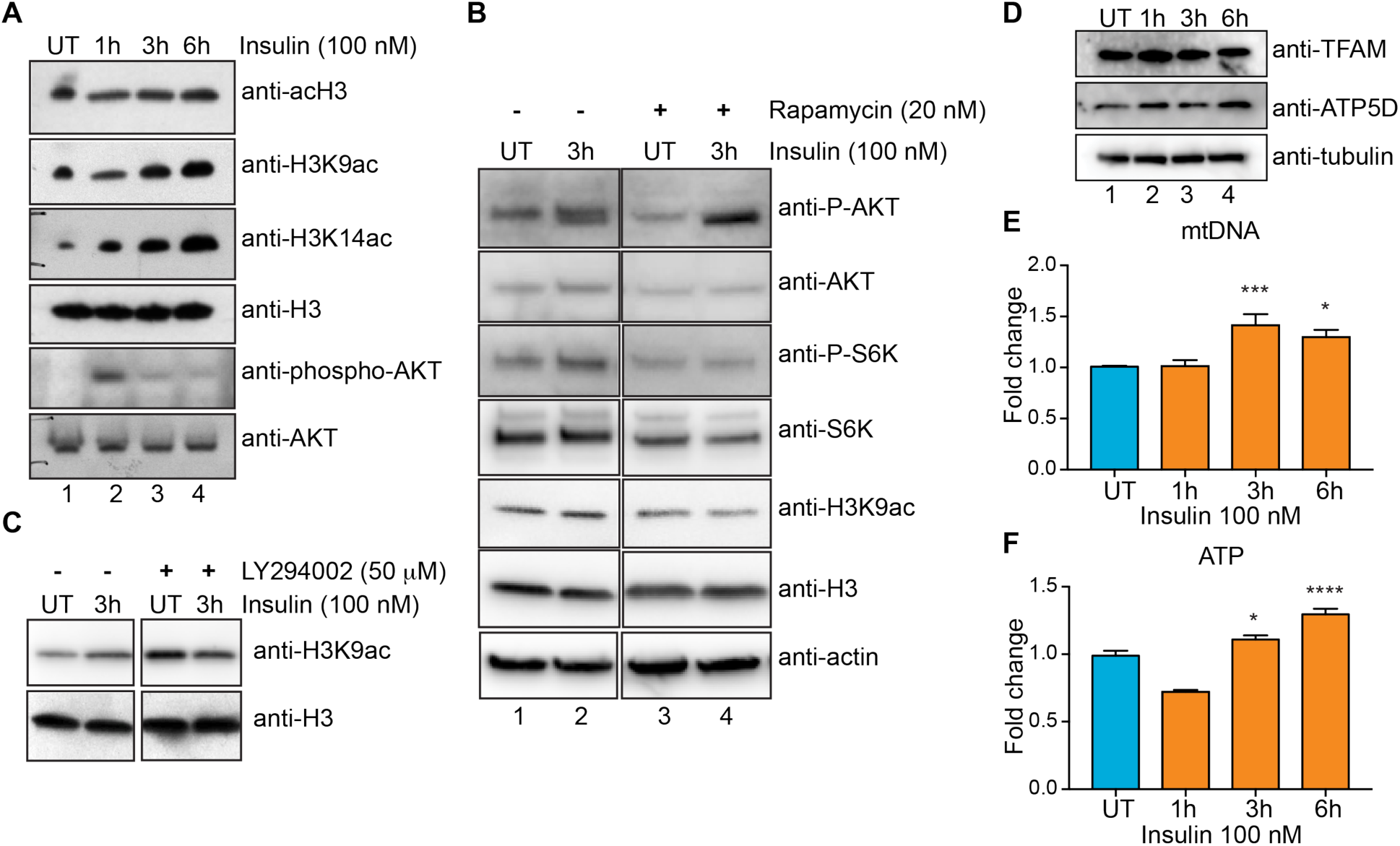
Insulin induces histone acetylation increases in MDA-MB-231 cells through PI3K-AKT-mTOR pathway and the mitochondria. (A) Western blot analysis using the indicated antibodies in MDA-MB-231 (TNBC cell line) cell lysates treated with insulin (100 nM) for 1h, 3h or 6h. (B) Western blot analysis using the indicated antibodies in MDA-MB-231 cells pretreated (lanes 3 and 4) or not (lanes 1 and 2) with 20 nM mTOR inhibitor rapamycin (1 h) followed by insulin treatment for 3h. (C) Western blot analysis using the indicated antibodies in MDA-MB-231 cells pretreated (lanes 3 and 4) or not (lanes 1 and 2) with 50 μM PI3K inhibitor LY294002 (1h) followed by insulin treatment for 3h. (D) Western blot analysis using the indicated antibodies in MDA-MB-231 cell lysates treated with insulin (100 nM) for 1h, 3h or 6h. (E) Bars show fold change in mitochondrial DNA content and (F) ATP levels in MDA-MB-231 cells treated with insulin (100 nM) for 1h, 3h or 6h. (E-F) Values are Mean+SEM from three independent experiments. Statistical significance was calculated using one-way ANOVA, Dunnett’s multiple comparisons test. *p<0.05, **p<0.01, ***p<0.001, ****p<0.0001.

mTOR complex 1 (mTORC1) enhances mitochondrial biogenesis and activity by promoting translation of nuclear encoded mitochondrial mRNAs including the components of Complex V and TFAM (transcription factor A, mitochondrial) (Morita et al., 2013). To test whether insulin induced histone acetylation increase was correlated with enhanced mitochondrial activity, we performed western blot analyses for TFAM and ATP5D (a subunit of the ATP synthase complex/Complex V) after insulin treatment. TFAM and ATP5D protein levels increase after 1h insulin treatment (Figure 1D). Next, we tested indicators of mitochondrial biogenesis and activity after insulin treatment. Mitochondrial DNA content is an indicator of mitochondrial number and ATP levels are a measure of mitochondrial activity in cells (Morita et al., 2013). Insulin increased the mitochondrial DNA content (Figure 1E) as well as ATP levels (Figure 1F) in MDA-MB-231 cells, indicating an enhancement in mitochondrial biogenesis and activity. In addition, we performed GC-MS to measure TCA cycle metabolites produced in the mitochondria. We observed increased levels of lactate and TCA cycle intermediates succinate, pyruvate, alpha-ketoglutarate, malate and citrate after 6h of insulin treatment (Figure S1C). These data suggest that insulin induces genome-wide increase in histone acetylation, in particular H3K9ac, through the PI3K-AKT-mTOR pathway.

### Insulin induces H3K9ac acetylation on promoter regions

To characterize the genomic loci associated with increased histone acetylation after insulin treatment, we performed quantitative ChIP-seq analyses as described in (Orlando et al., 2014) where we spiked in Drosophila S2 cells with MDA-MB-231 cells before the chromatin immunoprecipitation (ChIP) (see Methods). We observed an increase in the number of reads aligning to the human genome in the 3h and 6h insulin treated H3K9ac ChIPs indicating an increase in global histone acetylation levels (Table S1). By performing spike normalization (see Methods, Figure S2A-F), we observed global increases in H3K9ac levels on peaks in 3h and 6h insulin treated cells (Figure S2A-F).

Next, we annotated the H3K9ac peaks based on distance to the nearest RefSeq annotated TSS. Results showed that ~46% of the H3K9ac peaks were promoter proximal (~26% within 1kb and ~20% between 1kb-10kb of nearest TSS) (Figure 2A). We used DESeq2 (Love, Huber, & Anders, 2014) to identify peaks that were significantly increased (adjusted P<0.05) in both 3h and 6h insulin treated cells compared to UT. 22,372 and 9,171 peaks were significantly enriched with H3K9ac in 3h and 6h insulin treated cells respectively (Figure 2B and C). Of the significantly enriched peaks, about 58% and 55% of the peaks were promoter proximal (Figure 2A) in 3h and 6h samples respectively indicating that increases in H3K9ac were majorly localized at promoter regions and could potentially influence gene expression. Heat maps of H3K9ac at ±2kb around annotated start sites of transcripts further confirmed the increase in H3K9ac signal at promoter regions in insulin treated cells. There was a higher enrichment of H3K9ac signals at promoters in 3h compared to 6h treated cells (Figure S3). The highly enriched H3K9ac peaks had significant increases in H3K9ac signal at promoter regions (± 1kb from TSS) of their nearest genes (Figure 2D and E). These results show that insulin induces genome-wide increase in H3K9ac at promoter regions of genes and thereby could be involved in transcriptional regulation.

**Figure 2.**
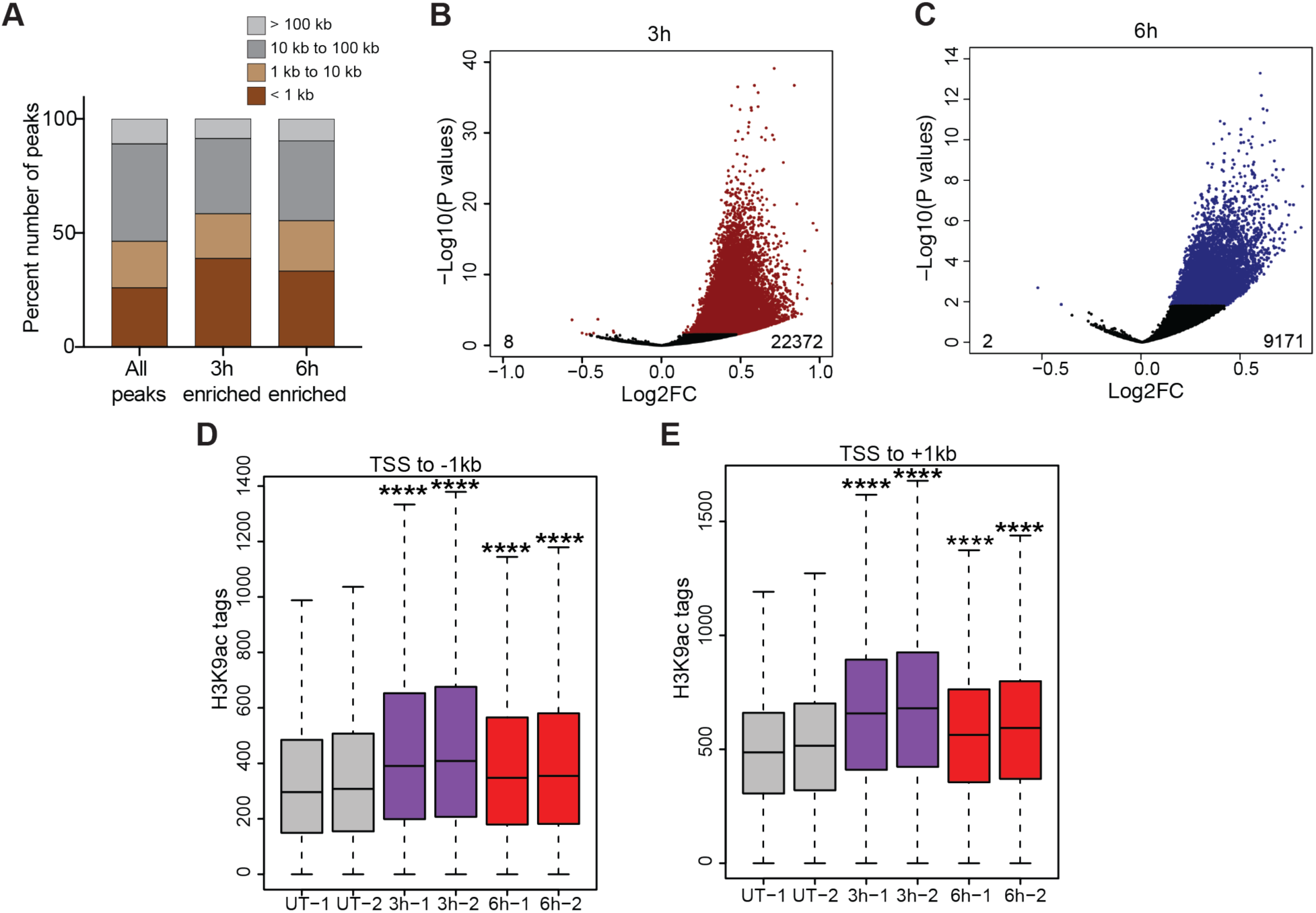
Insulin induces H3K9ac acetylation on promoter regions. (A) Stacked bars showing the distribution of H3K9ac peaks categorized by distance to nearest transcription start site (TSS). (B) Volcano plot showing the 22372 peaks that increased and 8 peaks that decreased H3K9ac acetylation after 3h insulin treatment. (C) Volcano plot showing the 9171 peaks that increased and 2 peaks that decreased H3K9ac acetylation after 6h insulin treatment. (D) Box plots showing the distribution of peak scores at -1kb to TSS regions of significantly enriched H3K9ac peaks. (E) Box plots showing the distribution of peak scores at TSS to +1kb regions of significantly enriched H3K9ac peaks. Significance was calculated using Kruskal Wallis test followed by Dunn’s multiple comparisons test. Adjusted p values were calculated using Benjamini-Hochberg method. ****p <0.0001.

### Insulin induces H3K9ac acetylation on promoters of insulin-induced genes

To further test whether the increase in H3K9ac enrichment levels correlate with gene expression changes induced by insulin, we performed RNA-sequencing (RNA-seq) in MDA-MB-231 cells untreated (UT) or treated for 3h and 6h with insulin. We quantified changes in gene expression after 3h and 6h insulin treatment from RNA-seq data using DESeq2 (Love et al., 2014). 207 and 384 genes exhibited significantly altered expression in in 3h and 6h insulin treated cells respectively (Figure 3A and 3B). Insulin treatment induced metabolic pathways required for cellular growth such as ribosome biogenesis, transcription, and splicing as well as known insulin regulated downstream pathways related to ATP production and mTOR signaling (Figure 3C). Insulin treatment downregulated FOXO signaling genes as well as apoptosis inducing genes. Interestingly, insulin treatment also downregulated genes involved in reactive oxygen species (ROS) metabolism or scavenging as well as immune cell migration and activation. Moreover, insulin upregulated several MYC (c-Myc) target genes and genes related to zinc ion homeostasis in cells (Figure 3C). These results indicate that insulin induces cell growth and proliferation while also suppressing apoptosis.

**Figure 3.**
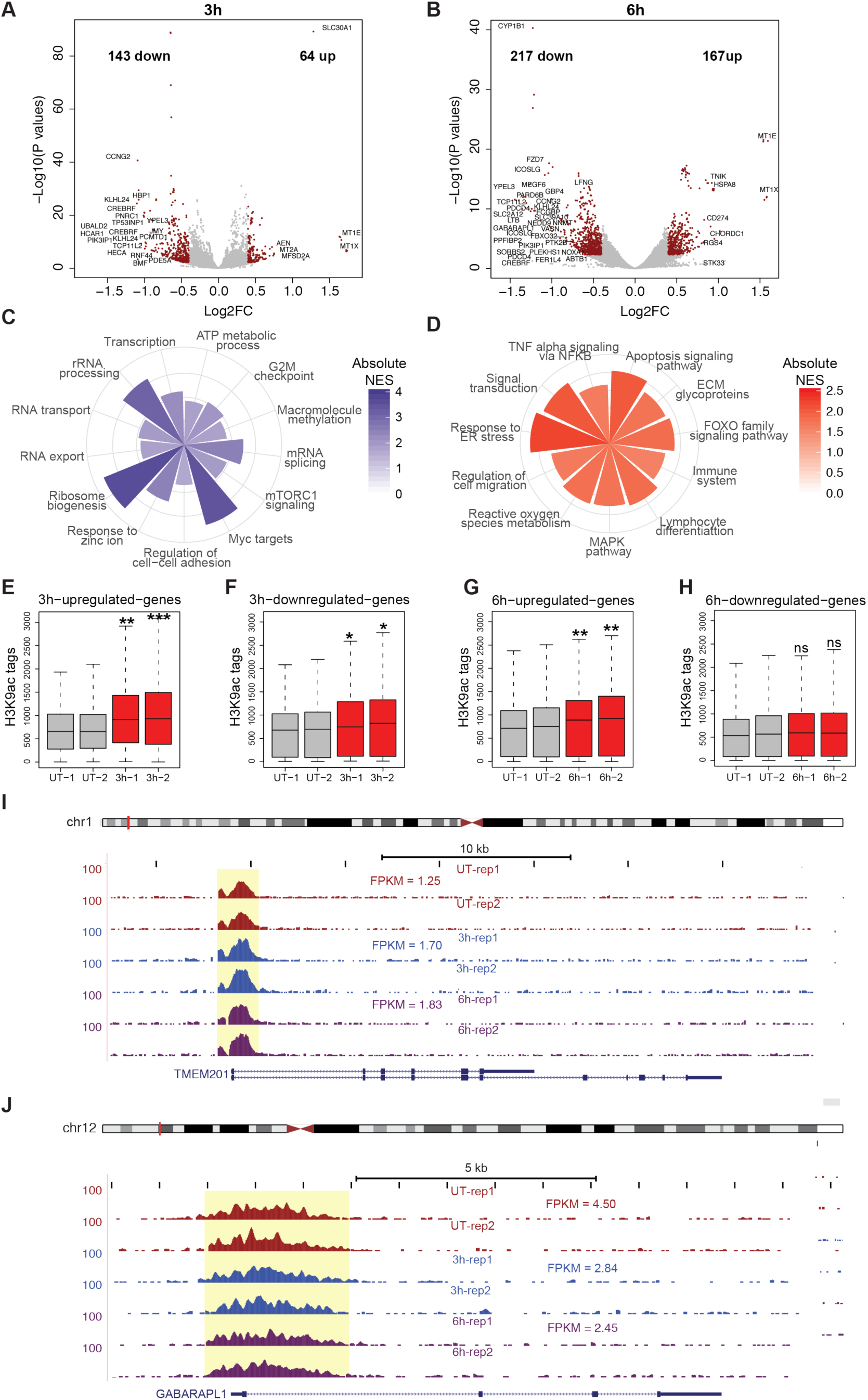
Insulin induces specific increases in H3K9ac acetylation on promoters of insulin induced genes. (A) Volcano plots showing the genes differentially expressed after 3h and (B) 6h insulin treatment. Differentially expressed genes are highlighted in red. (C) Gene sets enriched in Insulin upregulated genes and (D) Insulin downregulated genes. Absolute value of Normalized enrichment score (NES) from Gene Set Enrichment Analysis (GSEA) is shown. P < 0.05. (E-H) Box plots showing the normalized H3K9ac signal at promoters (TSS ± 1kb) of genes upregulated and downregulated after 3h or 6h of insulin treatment as indicated. Significance was calculated using Kruskal Wallis test followed by Dunn’s multiple comparisons test. Adjusted p values were calculated using Benjamini-Hochberg method. *p <0.05, **p <0.01, ***p <0.001, ****p <0.0001. (I) Genome browser screen shots showing the H3K9ac signal at *TMEM201* (upregulated gene) and (J) *GABARAPL1* (downregulated gene). Expression values (FPKM) are shown.

We then compared the changes in H3K9ac enrichment on the promoter regions (TSS ± 1kb) of genes upregulated and downregulated by insulin. Upregulated genes showed a larger increase in H3K9ac enrichment induced by insulin (Figure 3E and 3G) at 3h and 6h respectively. However, genes downregulated at 3h also showed a modest but significant increase in H3K9ac enrichment at their promoters (Figure 3F). Interestingly, genes downregulated after 6h insulin treatment did not show any significant enrichment in H3K9ac signals (Figure 3H). These results indicate that there is a genome-wide increase in H3K9ac signal at all expressed genes after 3h insulin treatment. However, at 6h the increase in H3K9ac signal is more specific to upregulated genes. Representative examples show the increase in H3K9ac enrichment on the upregulated gene *TMEM201* (Figure 3I) and no change in H3K9ac signal on the promoter of *GABARAPL1* (Figure 3J), a downregulated gene. These results indicate that the gene expression changes observed after insulin treatment are specific and are likely induced by signal-dependent transcription factors. Genome-wide increase in H3K9ac at promoters may facilitate increased chromatin accessibility at regulatory regions, however, is not enough to activate transcription which depend on recruitment of transcription factors, co-activators and RNA Polymerase II.

### Transcription factor NRF1 is involved in insulin mediated gene expression changes and histone acetylation

In order to further characterize the signal-dependent transcription factors involved in the altered gene expression network as well as chromatin acetylation induced by insulin, we performed transcription factor binding motif enrichment analyses. Genes upregulated by insulin showed a significant enrichment of E-box elements that are bound by transcription factors such as MYC (c-Myc), CLOCK, USF1, and BHLHE40 (Figure 4A). In addition to E-box elements, binding motifs for NRF1, ELF1, ELK1 and E2F transcription factors were also enriched. Interestingly, MYC target genes were also enriched in the upregulated gene set (Figure 3C), further implicating the involvement of MYC in enhancing expression of genes in response to insulin. Moreover, PI3K/AKT and MAPK pathways induced by insulin are known to enhance MYC activity by promoting the degradation of MAD1, an antagonist of MYC (Zhu, Blenis, & Yuan, 2008). We then categorized peaks with significantly increased H3K9ac signals based on distance from TSS of known RefSeq genes. We defined proximal peaks as peaks within 10kb of a known TSS and distal peaks as those outside this window. Proximal peaks with increased H3K9ac signals showed enrichment of transcription factor binding motifs for SP1, NRF1, ATF3, ELK1 and AP1 transcription factors among others (Figure 4B). Distal peaks that exhibited increased H3K9ac signal in response to insulin showed enrichment of FOS, JUN and AP1 family transcription factor binding sites (Figure 4C). NRF1 (Nuclear Respiratory factor 1) is required for expression of key metabolic genes regulating cellular growth. NRF1 also regulates the expression of several nuclear-encoded mitochondrial genes (Witkiewicz et al., 2011). As we earlier observed increased protein levels of mitochondrial proteins TFAM and ATP5D in response to insulin (Figure 1D), we tested whether NRF1 enrichment at its binding sites were enhanced after insulin treatment. ChIP experiments showed that NRF1 binding indeed increased after 3h insulin treatment at promoters of upregulated genes (Figure 4D-K). Increased NRF1 binding was also observed 6h post-insulin treatment, however, it was lower than that at 3h (Figure 4D-K) indicating an early response to insulin. NRF1 binding at promoters of NRF1 target genes could lead to increased histone acetylation at these regions. These results indicate that NRF1 might regulate the metabolic capacity of cancer cells by integrating metabolic inputs from the environment to increase histone acetylation on chromatin that allow continuous transcription from these genes.

**Figure 4.**
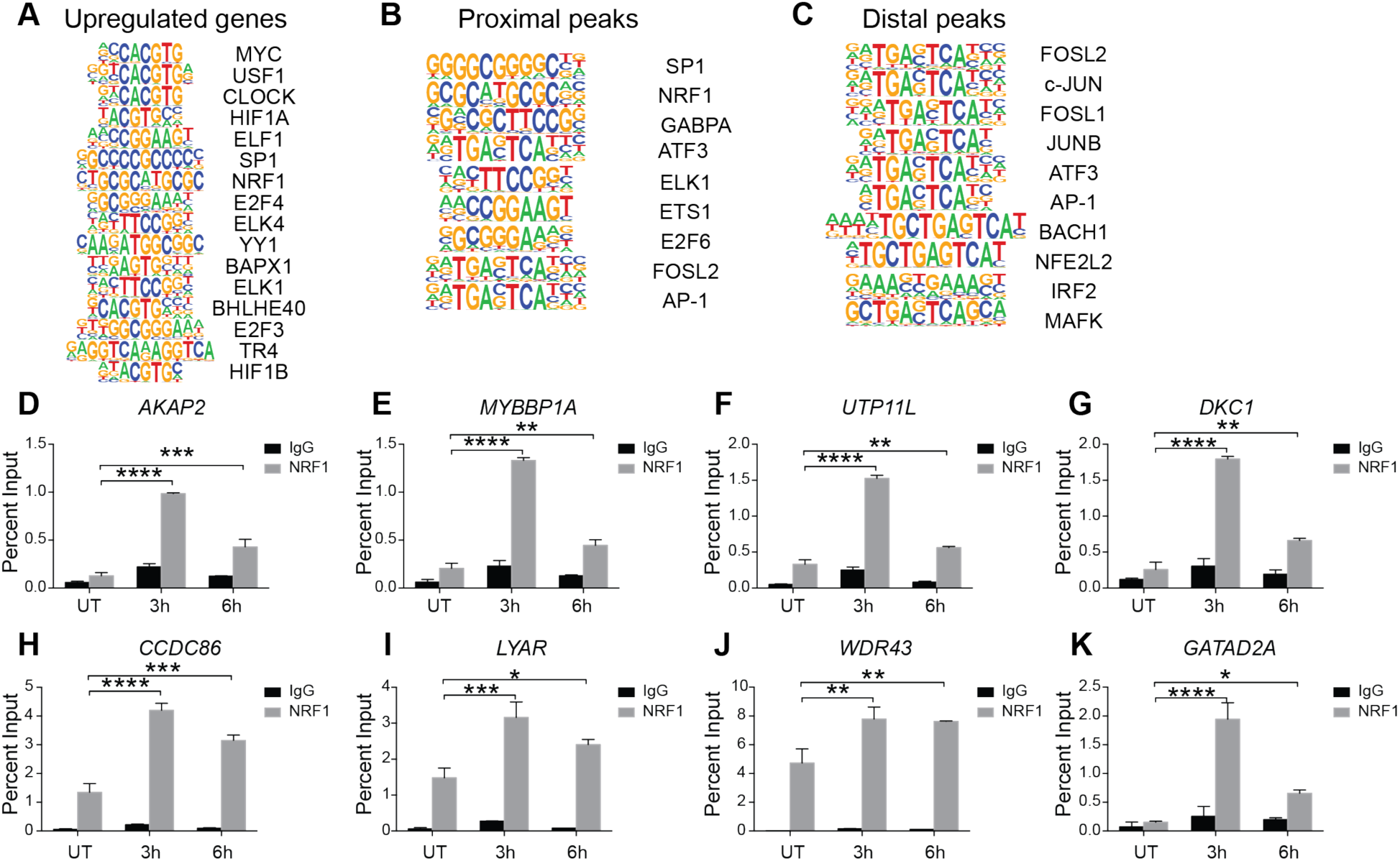
Transcription factors NRF1 is involved in insulin mediated gene expression changes and chromatin remodeling. (A) Transcription factor binding motifs enriched in promoter regions (TSS±1kb) of upregulated genes. (B) Transcription factor binding motifs enriched in promoter proximal and (C) distal H3K9ac peaks respectively. (D-K) NRF1 enrichment at NRF1 motifs in promoters of indicated insulin-upregulated genes determined by ChIP-qPCR in MDA-MB-231 cell lysates treated with insulin (100 nM) for 3h or 6h. Bars represent percent input pulldown in untreated (UT) and treated (3h, 6h) cells. Values are Mean+SEM from two independent experiments and three technical replicates from each experiment. Statistical significance was calculated using one-way ANOVA, Dunnett’s multiple comparisons test. *p<0.05, **p<0.01, ***p<0.001, ****p<0.0001.

### Insulin induced reactive oxygen species (ROS) causes genome instability

mTOR pathway induces mitochondrial biogenesis and activity that might lead to increased ROS production through the electron transport chain. To investigate whether insulin treatment induces ROS production in the cells, we measured ROS using a fluorescent dye, CellROX green. Results showed that ROS was significantly increased after 3h of insulin treatment and remained high at 6h (Figure 5A). Increase in ROS production could be deleterious to cells as the free radicals could cause DNA damage and mutation. We measured DNA damage using the DNA damage marker γ-H2AX in cells treated with insulin using immunofluorescence assays. We observed that the number of cells with γ-H2AX foci and the number of γ-H2AX foci per cell increased after 3h insulin treatment (Figure 5B). Interestingly, the number of cells with γ-H2AX foci decreased after 6h indicating possible activation of repair pathways (Figure 5B) as MDA-MB-231 cells harbor wild type *BRCA1*. To investigate whether hyperinsulinemia was associated with increase in histone acetylation and DNA damage in human samples, we measured the levels of H3K9ac and γ-H2AX in peripheral blood mononuclear cells (PBMCs) from an insulin-resistant and a healthy individual. We observed increased levels of H3K9ac and γ-H2AX in the insulin-resistant individual as compared to the insulin-sensitive individual (Figure 5C) corroborating our *in vitro* results. To investigate whether insulin induced chromatin changes can be reversed, we used metformin, a drug that is used for treatment of pre-diabetes and diabetes (Hostalek, Gwilt, & Hildemann, 2015). Metformin improves insulin sensitivity by several mechanisms, including mitochondrial Complex I inhibition and AMP kinase activation (Hur & Lee, 2015). Pre-treatment of cells with metformin prevented the insulin-induced increase in H3K9 acetylation (Figure 5D) indicating the potential significance of using metformin in triple negative breast cancer patients with hyperinsulinemia in preventing insulin mediated chromatin changes. Overall these data suggest that hyperinsulinemia might sensitize cells to DNA damage through increased ROS production and increased chromatin accessibility thereby potentially promoting deleterious mutations in pre-neoplastic lesions.

**Figure 5.**
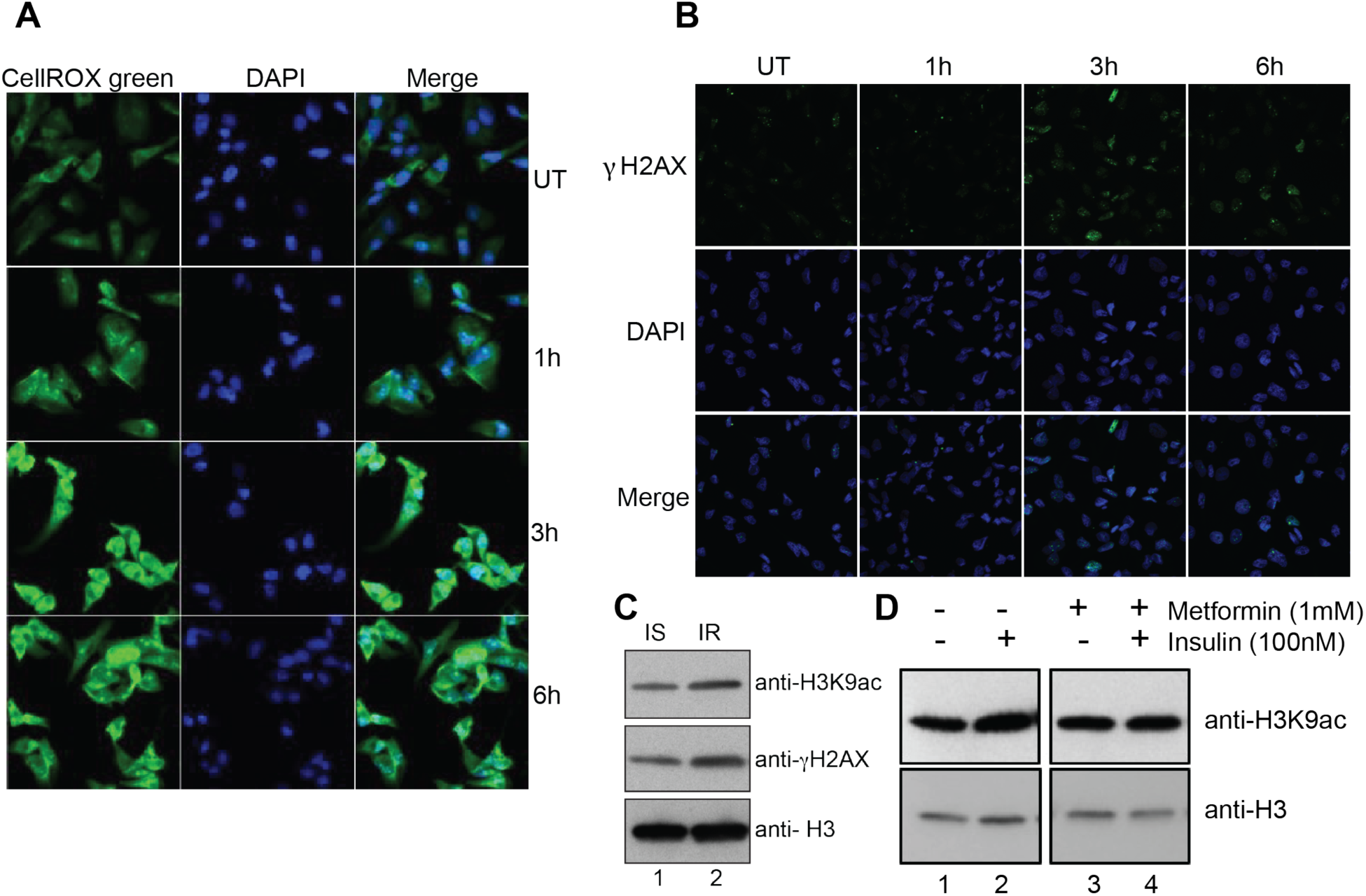
Insulin induced reactive oxygen species (ROS) causes genome instability. (A) ROS induction in insulin treated cells as measured by fluorescence signal from the CellROX green reporter. Nuclei are stained with DAPI. Magnification is 40X. (B) Representative immunofluorescence images showing intensity of γ-H2AX (DNA damage marker) (green) on MDA-MB-231 cells treated or untreated with Insulin (100 nM). Nuclei are stained with DAPI. Magnification is 20X. UT: Untreated (C) Western blot analysis for H3K9ac and γ-H2AX (DNA damage marker) on extracts from PBMCs isolated from insulin-sensitive (IS) and insulin-resistant (IR) individuals. (D) Western blot analysis using H3K9ac antibody in MDA-MB-231 cells pretreated (lanes 3 and 4) or not (lanes 1 and 2) with 1 mM metformin (24 h) followed by insulin treatment for 6h.

### Hyperinsulinemia enhances tumor growth in mice

To further demonstrate the role of hyperinsulinemia in enhanced tumor progression through altered chromatin acetylation, we used an immunodeficient hyperinsulinemic mouse model, Rag1^−/−^/MKR^+/+^ (Zelenko et al., 2016). MKR mice (Novosyadlyy et al., 2010) harbor a dominant negative mutation in the IGF-IR expressed specifically in the skeletal muscle. The female *Rag1*^−/−^MKR^+/+^ mice develop hyperinsulinemia but do not exhibit obesity, hyperglycemia or dyslipidemia. Orthotopic tumor xenografts were performed in *Rag1*^−/−^MKR^+/+^ (Rag/MKR) mice and *Rag1*^−/−^ (Rag/WT) female mice using MDA-MB-231 cells, as previously described (Shlomai et al., 2017). Tumors derived from the Rag/MKR mice were significantly larger and weighed more than those derived from the MKR tumors as shown earlier (Shlomai et al., 2017). To investigate whether tumors from hyperinsulinemic (MKR) showed increased histone acetylation, we performed western blot analysis on tumor protein extracts from Rag/WT or Rag/MKR mice. Results show a general trend towards increased histone acetylation in tumors from MKR mice (Figure 6C). To investigate the transcriptome changes associated with increased tumor growth in Rag/MKR mice, we performed RNA-seq analyses from two tumors each from Rag/WT and Rag/MKR mice. The MDA-MB-231 tumors derived from Rag/WT mice showed a completely different transcriptome compared to MDA-MB-231 cells grown in culture (data not shown) possibly due to differences in the microenvironment in the *in vitro* versus *in vivo* grown cells. The MDA-MB-231 tumors grown in Rag/MKR mice showed enhanced expression of genes related to collagen biosynthesis, extracellular matrix organization, cellular chemotaxis and interferon signaling (Figure 6D). Moreover, tumors grown under hyperinsulinemia showed decreased expression of FAS signaling, cell cycle and DNA damage checkpoint genes as well as genes that negatively regulate autophagy and PI3K-AKT signaling (Figure 6E). We also validated a few upregulated and downregulated genes from more tumors from each group by RT-qPCR to confirm the RNA-seq results (Figure 6F-I).

**Figure 6:**
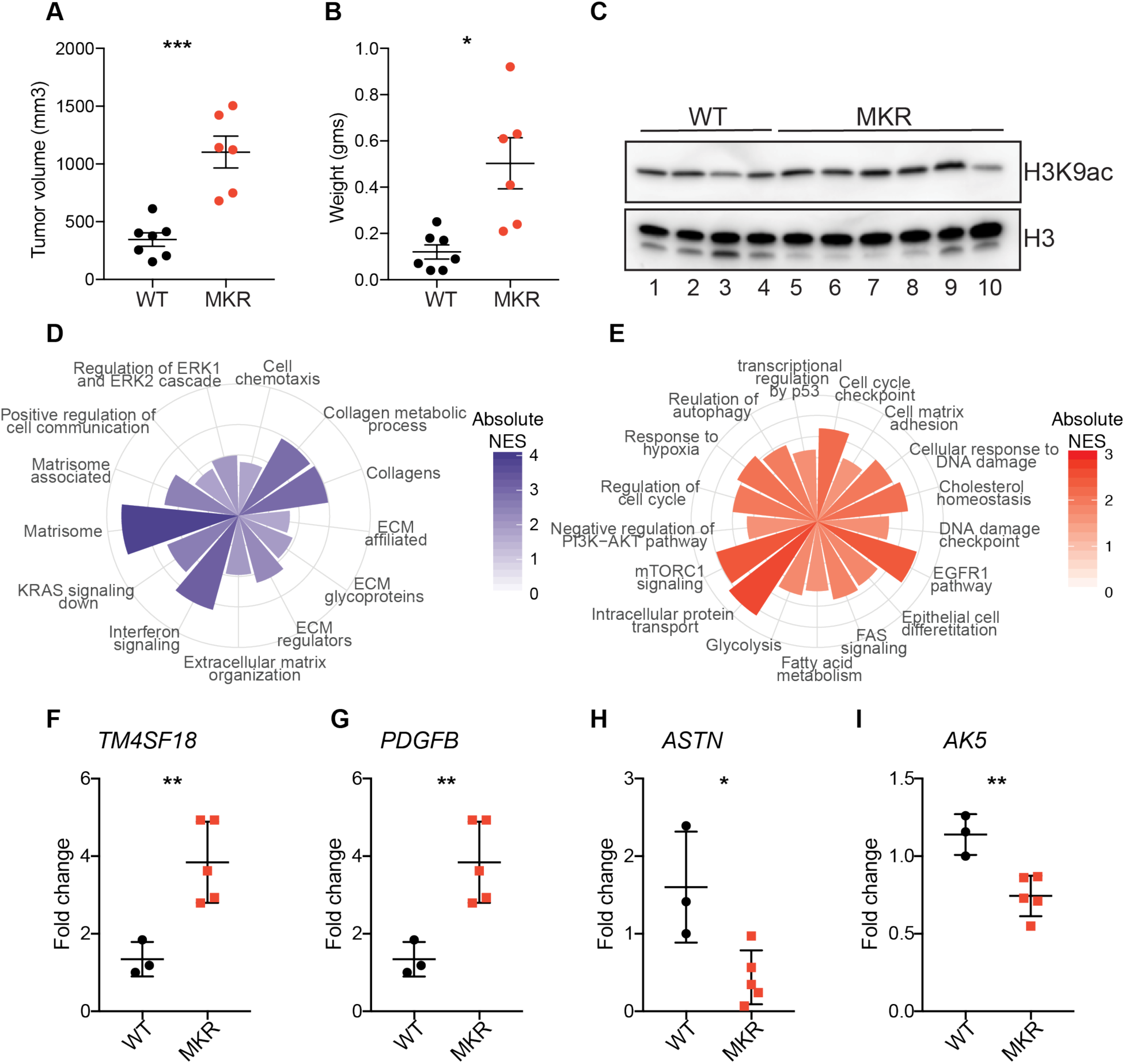
Hyperinsulinemia enhances ECM genes in mice tumors. (A) Tumor volume (mm^3^) and (B) weight were compared for tumors from Rag/WT (WT) and Rag/MKR (MKR) mice. Significance was calculated using unpaired Student’s t-test. *p <0.05, ***p <0.001. (C) Western blot analysis showing the total levels of H3K9ac and H3 in WT and MKR tumors. (D) Gene sets enriched in upregulated genes and (E) downregulated genes in Rag/MKR tumors compared to Rag/WT tumors. Absolute value of Normalized enrichment score (NES) from Gene Set Enrichment Analysis (GSEA) is shown. P < 0.05. (F-I) Gene expression quantification in WT and MKR derived tumors using RT-qPCR. Ct values were normalized with housekeeping gene 18S. Significance was calculated using unpaired Student’s t-test. *p <0.05, **p <0.01.

The gene expression changes induced by insulin in MDA-MB-231 cells cultured in vitro were not exactly mirrored by the gene expression changes in tumor xenografts in hyperinsulinemic mice. This observation is likely due to the fact that the MDA-MB-231 cells grown in vitro showed very different gene expression profiles as compared to the MDA-MB-231 tumors grown in Rag/WT mice. This is consistent with previous results demonstrating that breast cancer cells grown in a monolayer behave differently than those grown in a 3D microenvironment (Edmondson, Broglie, Adcock, & Yang, 2014). Moreover, the insulin treatment performed in *vitro* is a short (3h-6h) and acute (100 nM) exposure to insulin whereas tumors from Rag/MKR mice are exposed to chronic (several weeks) endogenous hyperinsulinemia. These factors may explain the differences in gene expression as well as pathways induced by insulin *in vitro* and *in vivo*. Altogether, our results show that hyperinsulinemia enhances MDA-MB-231 derived tumor growth possibly through enhanced histone acetylation.

## Discussion

Metabolic syndrome and its associated disorders are increasingly being recognized as enhanced risk factors for several types of cancers, including breast cancer. We investigated the impact of hyperinsulinemia, an important feature of metabolic syndrome, on chromatin and gene expression changes in TNBC cells. We observed genome-wide increases in chromatin-associated histone acetylation that was dependent on the insulin mediated signaling through the PI3K-AKT-mTOR pathway. We used a quantitative method of ChIP-seq (ChIP-Rx) to identify the regions associated with changes in H3K9ac in response to insulin. We found genome-wide increases in H3K9ac occupancy at gene promoters especially those that increased expression after insulin treatment. However, insulin-induced increase in histone acetylation at gene promoters was not always associated with an increase in gene expression indicating that increased acetylation at these sites may have a distinct function. Our observation is supported by a recent study investigating histone acetylation levels in response to high glucose levels (Lee et al., 2018). Interestingly, it has been proposed that histone acetylation may function as a capacitor for acetate/acetyl-CoA which could be utilized as an energy source or to balance the intracellular pH based on cellular condition (Kurdistani, 2014).

Insulin has long been proposed to be involved in breast tumor progression. Fifty percent of breast tumors and most established breast cancer cell lines including TNBC cell lines overexpress IR (Belfiore et al., 2009; Frasca et al., 1999; Gallagher & LeRoith, 2010; Law et al., 2008; Papa et al., 1990) and insulin induces proliferation in several breast cancer cell lines (Gliozzo et al., 1998). Our data further reinforce these observations. We find that insulin induces genes involved in ribosome biogenesis, transcription, splicing and metabolism that are regulated by MYC (c-Myc). Moreover, genes involved in apoptosis are downregulated in response to insulin.

Insulin enhances utilization of glucose by inducing mitochondrial activity and biogenesis through mTOR. Tumor cells are known to reprogram metabolic pathways to support the increased demand for macromolecules required for uncontrolled proliferation in response to growth factors. Acetyl-CoA is a central metabolite that links glucose metabolism to lipid synthesis as well as regulation of chromatin (Kinnaird, Zhao, Wellen, & Michelakis, 2016). Histone acetylation levels have been earlier shown to correlate with acetyl-CoA abundance (Cluntun et al., 2015). In proliferating cells, acetyl-CoA can be generated by a) oxidative decarboxylation of pyruvate from glycolysis; b) ATP-citrate lyase (ACLY) utilizing cytosolic citrate and c) ACSS2 using acetate (Kinnaird et al., 2016). ACLY activity is enhanced by AKT mediated phosphorylation thereby establishing a link between insulin signaling, acetyl-CoA levels and histone acetylation in the nucleus (Lee et al., 2014). The genome-wide increase in histone acetylation we observe in response to insulin might be a result of increased acetyl-CoA abundance due to increased ACLY activity.

Our findings also suggest that insulin induces ROS. This could be a result of increased mitochondrial activity as well as due to a decrease in scavenger proteins such as SOD2 (Superoxide dismutase 2). Indeed, we observe higher mitochondrial activity as well as decreased expression of SOD2 in response to insulin (RNA-seq results). Moreover, we find increased accumulation of DNA damage foci when the ROS levels peak after insulin treatment. These results indicate that increased insulin signaling might predispose cells to deleterious mutations if they fail to repair the damage. This might be very relevant in the context of BRCA1 mutated triple negative breast cancer cells.

We also show that tumors derived from hyperinsulinemic mice showed enhanced expression of extracellular matrix genes and deceased expression of DNA damage checkpoint genes which might help in tumor progression. However, the genes expressed in response to insulin exposure in the *in vitro* and *in vivo* conditions were markedly different, possibly due to the different microenvironments as well as interactions between tumor cells and extracellular matrix in the xenografted tumors. Given the link between hyperinsulinemia and the poor prognosis of breast cancer patients diagnosed with TNBC, understanding the cellular changes in response to hyperinsulinemia is important. Our finding that insulin drives hyperacetylation of histones in chromatin –thus impacting the transcriptome –highlights the impact insulin has within the nucleus.

## Materials and Methods

### Cell Culture

Human cell line MDA-MB-231 (Cat No. HTB-26) and *Drosophila* S2 cells (Cat No. CRL-1963) were purchased from American Type Culture Collection (ATCC, Manassas, VA, USA). MDA-MB-231 cells were grown in Dulbecco’s modified Eagle’s medium (DMEM) with high glucose (25mM) (Cat. No. 25-500; Genesee Scientific, San Diego, CA) at 37°C, 5% CO_2_ in a humidified chamber. S2 cells were cultured in Schneider’s *Drosophila* medium (Cat. No. 21720024; ThermoFisher Scientific, Waltham, MA) at 24°C in a humidified chamber without CO_2_. All media were supplemented with 10% heat-inactivated Fetal Bovine Serum (v/v) (SH30910.03, Fisher) and 1X antibiotics containing penicillin and streptomycin (Cat. No. 25-512, Genesee). Prior to insulin treatment, MDA-MB-231 cells were serum depleted in DMEM high glucose (25mM) medium containing 0.2% BSA (serum depletion medium) for 24h and then stimulated with 100nM insulin (Cat No. I9278; Sigma-Aldrich, St. Louis, MO) or left untreated (UT) for indicated time periods. Cell lines used in this study were tested and found to be mycoplasma negative.

### Western Blot

Cells were seeded at a density of 0.5 ×10^6^ cells per well in six well plates. The medium was changed to serum depletion medium 48h post-seeding. Insulin (100nM) treatment was done after 24h of serum depletion for indicated time durations. Cells were collected by scraping and centrifugation at 800×g for 5 mins at 4°C. Cell pellet was washed once with PBS and resuspended in 20 cell volumes of 1X SDS sample buffer. Cell lysates were prepared by performing two iterations of vortexing and heating at 95°C for 5 mins. Lastly, the cell lysates were sonicated using a Bioruptor^R^ pico (Diagenode, Leige, Belgium) for three cycles (30 sec on/30 sec off) at room temperature (RT) and cleared by centrifugation at 16000xg for 5 mins at RT. Western blotting was performed by running cell lysates on a gradient SDS-PAGE gel and transferring onto a PVDF membrane. Western blots were probed with antibodies specific to acetylated H3 (acH3) (Cat Nos. ab47915; Abcam, Cambridge, UK), H3K9ac (ab4441; Abcam), H3K14ac (C10010-1; EpiGentek, Farmingdale, NY), H3 (ab1791; Abcam), anti-phospho-AKT (2965S; Cell Signaling Technology, Danvers, MA), anti-AKT (2920S; Cell Signaling Technology), anti-phospho-S6K (9206S; Cell Signaling Technology), anti-S6K (9202S; Cell Signaling Technology), TFAM (7495S; Cell Signaling Technology), ATP5D (ab97491; Abcam), c-Myc (sc-40X; Santa Cruz Biotechnology, Dallas, TX), NRF1 (ab34682; Abcam), γH2AX (NB100-78356; Novus Biologicals, Littleton, CO), tubulin (2125S; Cell Signaling Technology) and actin (Sigma, A5441). After primary antibody incubation, blots were incubated with HRP-conjugated anti-rabbit or anti-mouse secondary antibodies (1:5000 dilution; Abcam, Cat. Nos. ab6721 and ab6789) and protein bands were visualized using chemiluminescence detection (Cat. No. 34076, Thermofisher Scientific).

### Mitochondrial DNA and ATP measurement

For mitochondrial DNA quantification, genomic DNA and mitochondrial DNA was extracted from insulin treated cells using DNeasy Blood and Tissue kit (Qiagen, Hilden, Germany). Genomic and mitochondrial DNA were quantified by qPCR using mitochondrial and genomic DNA specific primers; namely, cytochrome B and RPL13A (Table S2).

For ATP measurement, MDA-MB-231 cells untreated or treated with insulin in 12 well plates were lysed by boiling in deionized water for 5 mins. The lysate was clarified by centrifugation at 16000xg for 10mins at 4°C. ATP levels were measured using the luciferase-based ATP determination kit (Cat No. A22066; Thermofisher Scientific).

### Inhibitor treatment

MDA-MB-231 cells were pretreated with various signaling pathway inhibitors; LY294002 (50μM), MK-2206 (10μM), Rapamycin (20nM), Metformin (1mM) or vehicle DMSO for 1h prior to insulin treatment (100nM, 3h). The cells were then processed for western blot assays. LY294002 was purchased from Cell Signaling Technology (Cat No. 9901S). MK-2206 (Cat No. 11593) and Rapamycin (Cat No. 13346) were purchased from Cayman Chemical (Ann Arbor, MI). Metformin was purchased from Sigma-Aldrich (Cat No. D150959).

### Chromatin fractionation

Chromatin fractionation was performed using the method described in (Mendez & Stillman, 2000). Briefly, cells were collected from six well plates after insulin treatment by scraping and centrifugation at 200×g for 2 mins. Cell pellet was washed twice with PBS and resuspended and incubated in 100μl buffer A (10mM HEPES, pH 7.9, 10mM KCl, 1.5mM MgCl_2_, 0.34M sucrose, 10% glycerol, 1mM DTT) supplemented with protease inhibitors and 0.1% Triton X-100 for 8 mins on ice. Nuclei were isolated by centrifugation at 1300×g for 5 mins at 4°C. The nuclear pellet thus obtained was washed once with buffer A and then resuspended in 100μl buffer B (3mM EDTA, 0.2mM EGTA,1mM DTT) supplemented with protease inhibitors and incubated for 30 mins on ice. The nuclear suspension was centrifuged at 1700×g for 5 mins at 4°C to obtain nuclear soluble fraction in the supernatant and chromatin fraction in pellet. The chromatin fraction was resuspended in 1X SDS sample buffer and boiled at 95°C for 5 mins.

### Immunofluorescence analyses

MDA-MB-231 cells were grown on cover slips coated with poly-L-lysine at 37°C in a 5% CO_2_ incubator. After treatment, the DMEM medium was aspirated out and the cell layer was washed twice with PBS. The cells were fixed with 4% para-formaldehyde in PBS for 10 mins at RT. The para-formaldehyde solution was removed and quenched with 100mM Tris pH 7.2 for 5 mins at RT. The fixed cells were subsequently permeabilized by incubating in 0.1% Triton X-100 solution in PBS for 10 mins. The cells were then washed twice with PBS at 5 mins intervals and blocked using the blocking solution (10% FBS, 3% BSA in PBS containing 0.1% Triton X-100), for 45 mins at 37°C. After incubation, the blocking solution was replaced with anti-γH2AX antibody (Novus biologicals, Cat No. NB100-78356, 1 in 1000 dilution) for 30 mins at 37°C. The cells were then washed with the wash buffer (PBS containing 0.1% Triton X-100) twice for 5 mins each. Subsequently the cells were incubated with Alexa-488 conjugated anti-mouse antibody (Cat No. A-11029, 1 in 1000 dilution; ThermoFisher Scientific) for 30 mins at 37°C. The cells were washed with the wash buffer twice for 5 mins each.

For reactive oxygen species detection, insulin treated or untreated cells seeded on coverslips were treated with 5 μM CellROX Green reagent (Cat No. C10444; ThermoFisher Scientific) for 30 mins in culture medium at 37°C. Subsequently, the culture medium was aspirated and cells were washed thrice with PBS. Nuclei were stained using 10 μg/ml DAPI solution (Cat No. D1306; ThermoFisher Scientific) for 5 mins in the dark, followed by two washes with PBS. The coverslips were then inverted onto a microscopic slide over a drop of 70% Glycerol (in PBS) for visualization. The Alexa, DAPI and CellROX Green fluorescence were visualized with a Carl Zeiss confocal laser scanning microscope LSM 710 META. Images were captured using Zen software. ImageJ software was used to process the images. The images for comparative studies were captured at identical microscope settings.

### ChIP-Rx

ChIP-Rx was performed as described in (Orlando et al., 2014) with minor modifications. MDA-MB-231 cells were seeded at a density of 1 ×10^6^ cells in 60mm dishes and treated with 100nM insulin as described above. After treatment, cells were cross-linked using 1% formaldehyde for 10 mins at RT, followed by addition of 0.125 M glycine for 5 mins to stop the reaction. In parallel, S2 cells were crosslinked at a density of 1 ×10^6^ cells per ml using 1% formaldehyde. Crosslinked cells were then washed twice with ice-cold PBS supplemented with protease inhibitors. MDA-MB-231 and S2 cells were resuspended in parallel in cold lysis buffer 1 (140mM NaCl, 1mM EDTA, 50mM HEPES pH 7.5, 10% Glycerol, 0.5% NP-40, 0.25% Triton-X-100) supplemented with protease inhibitors (COmplete, Cat. No.11873580001, Sigma) and incubated on ice for 10 mins. Next, centrifugation was performed at 800×g for 5 mins at 4°C to pellet nuclei. Nuclear pellets were then resuspended in parallel in lysis buffer 2 (10mM Tris pH 8.0, 200mM NaCl, 1mM EDTA, 0.5mM EGTA) supplemented with protease inhibitors and incubated for 10 mins on ice. At this step, *Drosophila* nuclear suspension was added to each untreated and treated human MDA-MB-231 nuclear suspension at a ratio of (1 *Drosophila* cell per 2 human cells) or 1.5×10^6^ S2 cells to 3×10^6^ MDA-MB-231 cells. The composite cell nuclei were then pelleted at 800×g for 5 mins at 4°C.

Composite cell pellets were resuspended in SDS lysis buffer (1% SDS, 10mM EDTA, 50mM Tris-HCl, pH 8) supplemented with protease inhibitors and subjected to sonication using a Bioruptor^R^ pico for six cycles (30 sec on/30 sec off) to produce DNA fragments of 200–500 bp in length. Sheared chromatin was clarified by centrifugation at 16000×g for 5mins at 4°C. 0.3 ×10^6^ cell equivalents were diluted with ten volumes of cold ChIP-dilution buffer (0.01% SDS, 1.1% Triton X-100, 1.2 mM EDTA, 16.7 mM Tris-HCl, pH 8, 167 mM NaCl) supplemented with protease inhibitors and used for each immunoprecipitation. About 10% of the chromatin from each ChIP reaction was saved as input. ChIP assays were performed with 5μg anti-H3K9ac antibody and 25μl of magnetic protein G Dynabeads (Cat. No. 10004D; Thermofisher Scientific) which were incubated overnight at 4°C. 5μg rabbit IgG was used for control ChIPs. Magnetic beads were washed successively with low salt buffer (0.1% SDS, 1% Triton X-100, 2mM EDTA, 20mM Tris-HCl, pH 8, 150 mM NaCl), high salt buffer (0.1% SDS, 1% Triton X-100, 2mM EDTA, 20mM Tris-HCl pH 8, 500mM NaCl), LiCl buffer (250 mM LiCl, 1% NP40, 1% NaDOC, 1mM EDTA, 10mM Tris-HCl, pH 8), and twice with TE buffer (10mM Tris-HCl pH 8, 1mM EDTA). Elution buffer (1% SDS and 100mM NaHCO_3_) was added to the washed beads, and the bead solution was incubated at RT for 30 mins. In parallel, the saved input was also diluted in Elution buffer. The DNA-protein complexes were then reverse cross-linked by adding 200mM NaCl, 20 μg Proteinase K (Cat. No. P4850; Sigma-Aldrich) and incubating at 65°C for 4 hours. Subsequently, 20 μg of RNase A (Cat No. EN0531; ThermoFisher Scientific) was added and further incubated for 15 mins at 37°C. The immunoprecipitated DNA was extracted using phenol-chloroform and ethanol precipitation. Resultant ChIP DNA was quantified using Quant-iT™ dsDNA Assay Kit (Cat No. Q33120; ThermoFisher Scientific) and used for library preparation.

ChIP-seq libraries were made using Illumina Tru-Seq library preparation kit (Illumina, San Diego, CA) and multiplexing barcodes compatible with Illumina HiSeq 2500 technology. About 50 million single-end reads of length 51 bp were generated from each ChIP-seq library.

### ChIP-seq analyses

Sequencing reads from each library were aligned to a combined reference genome (human + *Drosophila*) using bowtie (Langmead, Trapnell, Pop, & Salzberg, 2009). The combined reference genome was generated as described in (Orlando et al., 2014). Briefly, a combined genome sequence was created by concatenating the genome sequences of human (hg19) and *Drosophila* (dm3). Next, custom bowtie indexes were generated for the combined genome sequence using ‘bowtie-build’ command. Bowtie alignment was done against the combined genome using parameters: -m 1 -e 70 -k 1 -n 2 --best --chunkmbs 200. About 6% of reads aligned to the dm3 genome and ~94% aligned to the hg19 genome. The number of reads aligned to human and *Drosophila* genome are reported in Table S1. We identified a union set of 40,222 and 5,716 H3K9ac peaks in the human and *Drosophila* cells respectively. We normalized peak scores for the 40,222 human (hg19) peaks using hg19 aligned read counts (read count normalization) (Figure S2A). Moreover, we used *Drosophila* (dm3) aligned read counts for normalizing peak scores (spike normalization) (Figure S2B). Significantly, we observed greater changes in H3K9ac levels on peaks in 3h and 6h insulin treated cells after spike normalization as compared to read count normalization (Figure S2C and D). The effect of spike normalization was also evident in aggregate profiles of H3K9ac ±2kb around annotated transcription start sites (TSSs) (Figure S2E and F). Overall spike normalization led to better conformity between replicates and revealed global increase in histone acetylation that could be quantified.

### RNA-seq analyses

Total RNA was isolated from insulin treated cells or from tumors tissues using NucleoSpin^®^ RNA kit (Macherey-Nagel, Germany) with on-column DNase I digestion. PolyA-enriched RNA was isolated and used for library preparation using TruSeq RNA library Prep kit (Illumina). About 50 million single-end reads of length 51 bp were generated for the MDA-MB-231 samples and 50 million paired-end 100 bp reads were generated from the MDA-MB-231 xenograft tumors. Raw sequences were aligned to the hg19 reference genome using HISAT2 2.1.0 (Kim, Langmead, & Salzberg, 2015) using default parameters. Stringtie 1.3.4 (Pertea et al., 2015) was used with default parameters to assemble transcripts using the Genocde v19 transcript annotation. Assembled transcripts from all libraries were further merged using --merge option in Stringtie. Merged transcript abundances were measured using bedtools coverage and DESeq2 package (Love et al., 2014) was used to normalize counts and identify differentially expressed genes (log2 fold change ≥ 0.5 and padj < 0.1). Gene Set Enrichment Analysis (GSEA) (Subramanian et al., 2005) was used to determine significantly altered gene ontology and pathways. For validation of RNA-seq data, 1μg total RNA was used to synthesize cDNA using High-Capacity cDNA Reverse Transcription Kit (Cat No. 4368814; ThermoFisher Scientific). Gene expression was analyzed by quantitative PCR (qPCR) using KAPA SYBR^®^ Fast ROX Low qPCR Master (Cat No. KM4117, Kapa Biosystems, Inc., Wilmington, MA) using gene-specific primers (Table S2). Relative gene expression between groups was determined using 2^^-ΔΔCt^ method after normalization with 18S levels.

### Visualization of ChIP-seq data and additional analyses

Reads aligning to the hg19 genome were filtered out from the aligned bam files and peaks were called for each library with respective Input using macs2 (Zhang et al., 2008) with broad peak calling option and q < 0.1 for broad regions and q < 0.05 for narrow regions. Peaks were annotated using annotatePeaks.pl command in homer (Heinz et al., 2010) and assigned to the nearest hg19 RefSeq gene TSS. Wiggle tracks were generated using custom scripts, normalized by the number of dm3 aligned reads and visualized on the UCSC Genome Browser. Heatmaps were generated using Java TreeView (Saldanha, 2004) and aggregate profiles were made using deepTools (Ramirez et al., 2016). Motif enrichment analysis was performed using findMotifsGenome.pl command in homer.

### Human samples

Blood samples were obtained from patients after an 8h fasting period following institutional guidelines at the City of Hope (IRB no. 15418) in purple-top EDTA vacutainer tubes after obtaining written informed consent. Insulin resistance was identified by measuring Hemoglobin A1C (HbA1c) using HPLC method (Davis, McDonald, & Jarett, 1978). HbA1C levels between 5.7–6.3 were used to define insulin-resistance. All individuals were within ages of 18–55. PBMCs were isolated from whole blood using Ficoll-Paque method. Briefly, whole blood was diluted 1:1 with PBS containing 2% FBS, layered on top of 15 ml Ficoll and spun down at 1200xg for 10 mins. The white buffy coat containing PBMCs were collected and washed twice with PBS containing 2% FBS and spun down at 1200xg for 10 mins to remove platelets. 0.5 x10^6^ PBMCs were lysed in 20 cell volumes of 1X SDS sample buffer and processed for western blot analyses.

### Tumor xenograft studies

Animal studies were performed at the Icahn School of Medicine at Mount Sinai (ISMMS) Center for Comparative Medicine and Surgery. All studies were approved by the ISMMS Institutional Animal Care and Use Committee. All animals used for the studies, were female, on an FVB/n background. The immunodeficient hyperinsulinemic mice were generated as previously described by crossing the recombination-activating gene 1 (*Rag1*) knockout mice with the muscle creatinine kinase promoter expressing dominant-negative *Igf1r* (MKR) mice (Zelenko et al., 2016). The metabolic phenotyping of the Rag1 knockout (Rag1^−/−^) / MKR mice and control Rag1^−/−^ mice has been characterized previously (Zelenko et al., 2016). Mice for this study were maintained on regular diet (PicoLab Rodent Diet 20, 5053), with free access to water and a 12 hour light / dark cycle. 5×10^6^ MDA-MB-231 tumor cells were injected into the inguinal mammary fat pad of *Rag1^−/−^* and Rag*^−/−^*/MKR female mice aged between 8 and 10 weeks. MDA-MB-231 tumor growth was measured as previously described (Shlomai et al., 2017). At the end of the study, tumors were dissected and flash frozen in liquid nitrogen for further analysis.

### Statistical analyses

Data are represented as mean and standard error of mean (Mean + SEM). Statistical analyses were performed using GraphPad Prism 7.0 software (GraphPad Prism Software Inc., San Diego, CA) and R. Normal distribution was confirmed using Shapiro-Wilk normality test before performing statistical analyses. For normally distributed data, comparison between two means were assessed by unpaired two-tailed Student’s t test and that between three or more groups were evaluated using one-way analysis of variance (ANOVA) followed by Tukey’s post hoc test. In case of Student’s t-test, F-test was performed to check whether the variance in the groups compared were significantly different. For data where significantly different variances were observed, t-test with Welch correction was performed. For data that did not follow a normal distribution, Mann-Whitney test was performed for comparison between two groups and Kruskal Wallis test followed by Dunn’s multiple comparisons test was performed for comparing more than two groups. A p-value of < 0.05 was considered statistically significant. Figures were generated using Adobe Illustrator software (San Jose, CA, USA).

## Acknowledgments

Research reported in this publication was supported by the City of Hope CCSG Pilot award from National Cancer Institute of the National Institutes of Health under award number P30CA033572 and the National Cancer Institute grant (Grant Number: R01CA220693). We acknowledge services provided by the Integrative Genomics, Light Microscopy and DNA/RNA synthesis cores supported by the National Cancer Institute of the National Institutes of Health under award number P30CA033572. The content is solely the responsibility of the authors and does not necessarily represent the official views of the National Institutes of Health.

## Competing Interests

The authors declare that no competing interests exist.

