## Supplemental data for "Hyperinsulinemia promotes aberrant histone acetylation in triple negative breast cancer"

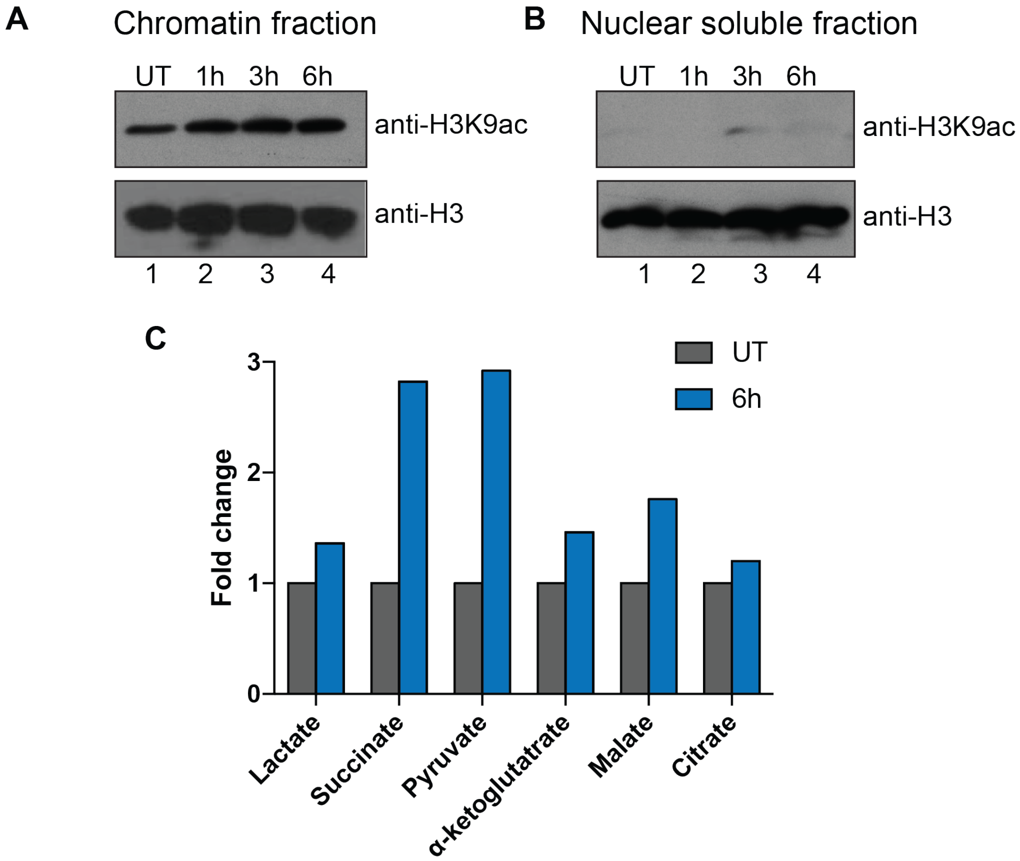


**Figure S1:** (A) Western blot analysis for H3K9ac in chromatin fraction (left panel) or (B) nuclear soluble fraction (right panel) extracted from insulin treated cells. (C) Bars represent fold change in levels of indicated metabolites in cells treated with 100nM insulin for 6h. UT: Untreated.

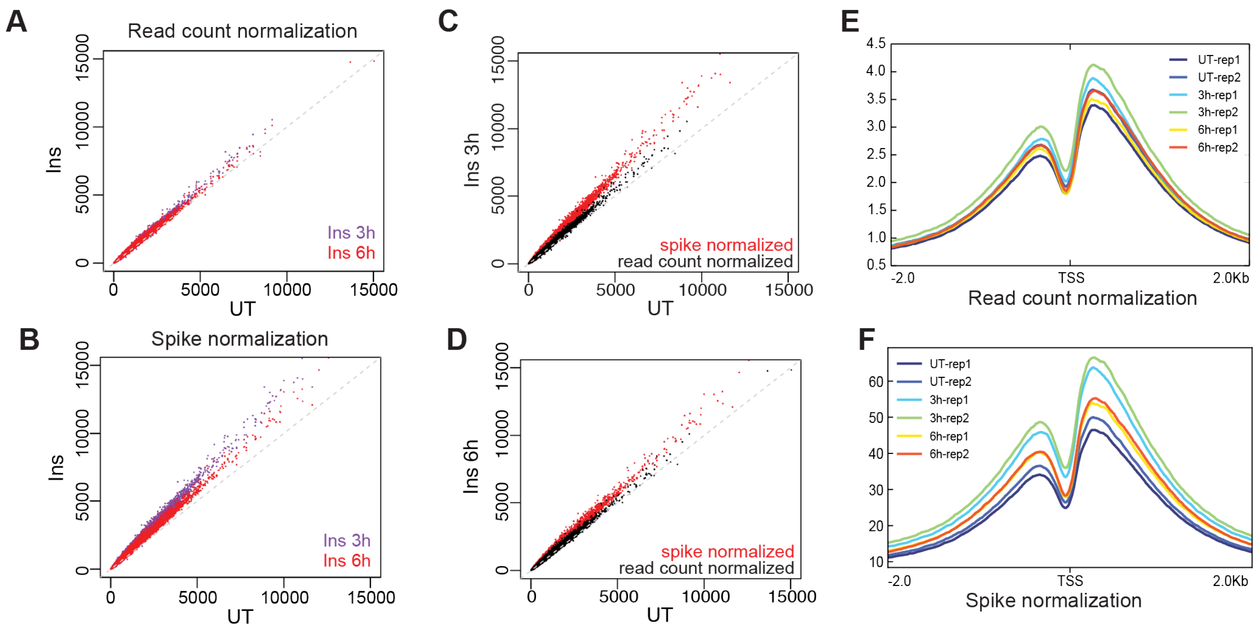


**Figure S2: Spike normalization quantifies changes in histone acetylation.** (A) Scatterplots showing the peak scores in UT versus 3h (purple) and 6h (red) Insulin treatment normalized with total aligned read counts (hg19) and (B) spike-in normalization (aligned read counts from dm3 genome). (C) Scatterplots showing the peak scores in UT versus 3h and (D) UT vs 6h Insulin treatment normalized with total aligned read counts (hg19) (black) or spike-in normalization (red). (E) Aggregate profile of H3K9ac signals around transcription start sites (TSS) normalized using read count normalization or (F) spike normalization in all ChIP-seq libraries.

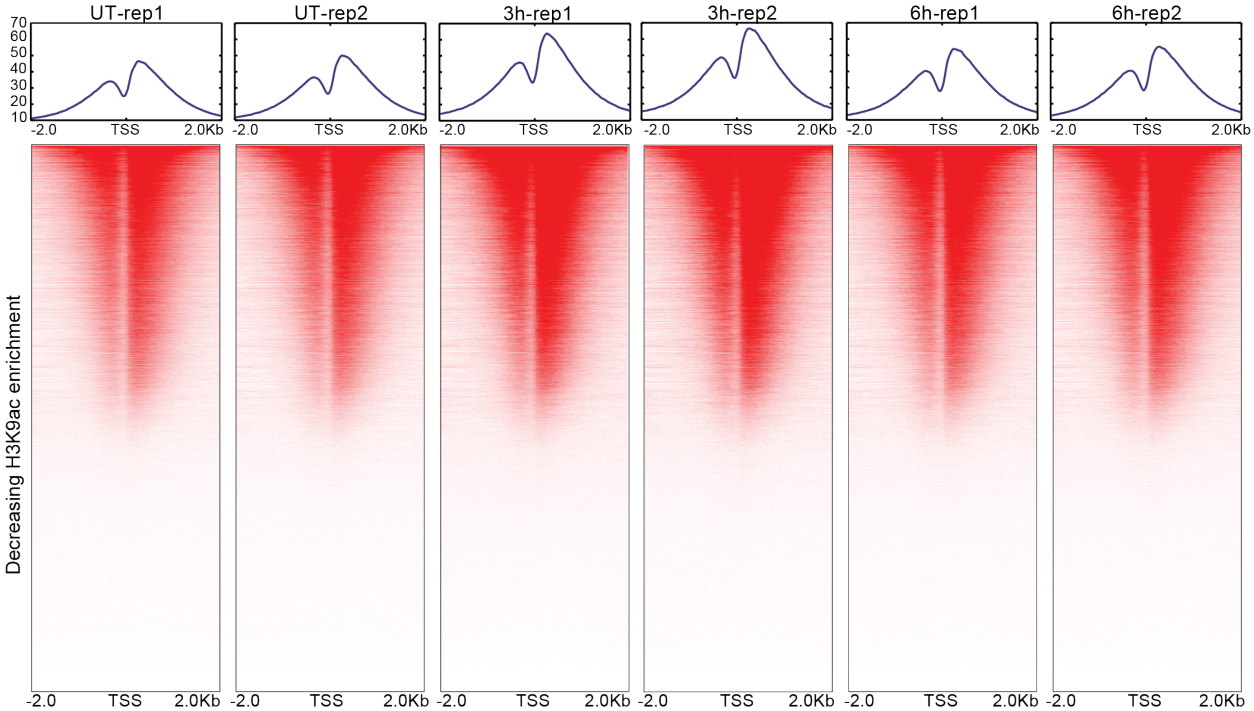


**Figure S3:** Heat maps and average profiles showing the distribution and enrichment of H3K9ac tags at promoter regions (±2kb from TSS of all transcripts in UT and 3h and 6h Insulin treated samples.

### Supplementary Tables

| Sample | Number of reads | uniquely aligned reads | uniquely aligned reads after PCR duplicate removal | hg19 aligned reads | dm3 aligned reads |
|---|---|---|---|---|---|
| UT-rep1 | 53085048 | 43556151 | 40433495 | 37702767 | 2730728 |
| UT-rep2 | 48979747 | 39584625 | 36880664 | 34371563 | 2509101 |
| 3h-rep1 | 51726037 | 42828559 | 39670425 | 37411094 | 2259331 |
| 3h-rep2 | 46332700 | 37745757 | 35166227 | 33128942 | 2037285 |
| 6h-rep1 | 52300364 | 42586950 | 39937204 | 37518909 | 2418295 |
| 6h-rep2 | 56543804 | 45720199 | 42239894 | 39637418 | 2602476 |
| UT-Input | 52847588 | 41039624 | 39296328 | 37155271 | 2141057 |
| 3h-Input | 57199048 | 45243608 | 42952024 | 41059938 | 1892086 |
| 6h-Input | 53845368 | 41840849 | 39643942 | 37586258 | 2057684 |

**Table S1: Read number information**

Table shows the number of sequencing reads obtained, uniquely aligned reads before and after PCR duplicate removal, number of reads aligned to hg19 (human) genome and dm3 (*Drosophila*) genome for each library.

| **Genomic DNA and mitochondrial DNA primers** | | |
|---|---|---|
| **Gene** | **Forward primer (5'-3')** | **Reverse primer (5'-3')** |
| *MT-CYB* (cytochrome B) | gcgtccttgccctattactatc | cttactggttgtcctccgattc |
| *RPL13A* | cttgctggtcttcgttcaaatc | gaggaacagggactgagaaag |
| **RT-qPCR primers** | | |
| **Gene** | **Forward primer (5'-3')** | **Reverse primer (5'-3')** |
| *TM4SF18* | catctctgccttgggtcttg | ccactgagggttatgagaatgg |

| | | |
|---|---|---|
| *PDGFB* | agtcggcatgaatcgctg | tcatgttcaggtccaactcg |
| *ASTN* | ggcgatgtgaggatgagttag | gtagggtttgatctcgctgtag |
| *AK5* | aacgatgccaaggagtatctg | gtattctactggatcttcgggc |
| **ChIP primers** | | |
| **Gene** | **Forward primer (5'-3')** | **Reverse primer (5'-3')** |
| *DKC1* | ttccagcctgggccaac | ctggtcgtctgcgcaat |
| *CCDC86* | AAAGACTGGCTCATCAATC ACA | CGGCCATGTTGGTGAGG |
| *AKAP2* | CGGCTCCTGGCATTTGA | TCGCTGTGTGTGCATGT |
| *LYAR* | CGCAGCTACCTGCCTCT | GAACCGCCTTCCTGCTTC |
| *WDR43* | TCCTGCGACGCGAAGAT | CCGCCATTGCTGCTCTG |
| *GATAD2A* | GAGACTGAGCCGCGAGA | GACAGACGACCGACCGA |
| *MYBBP1A* | AATTACATTGAACTCATCT GACTGG | AGCTGCCGACTGCATTAT |
| *UTP11L* | TACCTTCTAGATACAGCAA CCC | CTACCCAGATCTCGCCTTC |

**Table S2: Sequences of primers used in the study**
